# PAM-interacting domain swapping is extensively utilized in nature to evolve CRISPR-Cas9 nucleases with altered PAM specificities

**DOI:** 10.1101/2021.05.01.442224

**Authors:** Qiuyan Wang, Xiaomeng Bian, Jinli Du, Yawen Lv, Lin Tao, Tian Xie

## Abstract

It has been anticipated that protospacer adjacent motif (PAM) specificity of the CRISPR-Cas9 nucleases protospacer appears to be modular. Here we present the finding that naturally occurring domain swapping has been extensively involved in altering its PAM specificities. Sequence analysis of streptococcal Cas9 sequences revealed conservation of three distinct PAM-interacting motifs, with phylogenetic analysis of full-length Cas9 and the PID demonstrating that PAM-interacting domain (PID) domain swapping was extensively utilized to diversify its PAM specificity. An extended analysis of 582 representative Cas9 sequences revealed that this PID swapping was broadly present in most of the investigated genera. Mimicking the natural PID domain swapping, a functional chimeric enzyme, based on the scaffold of compact Staphylococcus aureus Cas9, with novel NNAAAA PAM specificity was developed. In summary, our findings shed new light on a rich source of exchangeable PID domains in Cas9 genes, which can be mined for domain swapping aiming to an effective PAM refinement.

## Introduction

The CRISPR/Cas9 system was originally discovered in prokaryotic organisms as an adaptive immunity mechanism used to cleave invading nucleic acids (1–3). CRISPR-Cas9 nucleases enable efficient genome editing in a wide variety of organisms and cell types (1). Target-site recognition by Cas9 is directed by CRISPR RNA (crRNA) and trans-activating crRNA (4) and requires recognition of a short neighboring protospacer adjacent motif (PAM). Because Cas9 systems are easier and cheaper to implement than other comparable technologies, CRISPR technology has been wildly utilized for applications in genome editing, gene regulation, and DNA imaging and are being explored for gene drives and sequence-specific antimicrobials (5–8). However, the broad utility of Cas9 in numerous applications is restricted by its PAM compatibility, as the strict PAM-reorganization pre-requirement hinders applications of the CRISPR-Cas9 system due to the inaccessibility of genetic loci lacking PAMs.

To relax this constraint, additional Cas9 variants with distinct PAM requirements are urgently needed. Several natural CRISPR nucleases with different PAM requirements have been developed as alternative tools, including *Staphylococcus aureus* Cas9 (9), *Neisseria meningitidis* Cas9 (10), *Francisella novicida* Cas9 (11), *Campylobacter jejuni* Cas9 (12), and *Acidaminococcus* sp. Cpf1 (13). Additionally, rational design and irrational design have been applied to refine Cas9 PAM specificity. For example, the VQR (D1135V/R1335Q/T1337R), EQR (D1135E/R1335Q/T1337R), and VRER (D1135V/G1218R/R1335E/T1337R) variants of *Streptococcus pyogenes* Cas9 (SpCas9), which recognize 5’-NGAN-3’, 5’-NGNG-3’, and 5’-NGCG-3’ PAMs, respectively, were developed by directed evolution (14). Moreover, SpCas9-NG was rationally engineered to recognize relaxed 5’-NG-3’ PAMs by introducing non-base-specific interactions with the PAM duplex to compensate for the loss of base-specific interactions (15). Based on SpCas9 structural information and the amino acid–DNA-interacting information obtained from SpCas9 variants, a structurally engineered and nearly PAM-less SpCas9 variant (SpRY) was developed to be capable of targeting most DNA sequences with high editing efficiency and flexibility (16). Although, serial successful PAM relaxed SpCas9s were created, the rational design and irrational design of PAM for Cas9 remain intricately difficult tasks. For rational design, the complexity of interactions between binding sites and PAM makes predictions difficult. Previous attempts to change SpCas9 PAM specificity to an NAA PAM by simply introducing arginine-to-glutamine substitutions at positions 1333 and 1335 were unsuccessful, thereby ruling out a simple PAM-recognition code (17). Irrational design of PAM alterations is labor-intensive and time-consuming, as validation requires high-throughput PAM-determination assays. Additionally, random mutagenesis with a significant bias for transitions over transversions and absence deletion and insertion will limit access to a broader sequence space (18). Therefore, customized PAM engineering remains highly challenging, and alternative strategies for generating Cas9 variants with new PAM specificities are urgently needed.

PAM-interacting domain (PID) shuffling might be an appealing alternative, given that it is efficient at harnessing the natural diversity. In this study, we determined that naturally occurring PID swapping has been broadly utilized in the Cas9 family to provide rapid PAM adaptation to race with rapidly evolving phages. Inspired by this finding, we suggest that PID swapping should in principle be extendable to a wide variety of Cas9 orthologues in order to take advantage of the natural diversity of PAM specificity.

## Results

### Detection of Cas9 homologs with PID domain swapping

Sequence-specific PAMs harboring Arg1333 and Arg1335 at Cas9 positions at interaction sites with the DNA duplex (D1332RKRY1336) were well conserved in other type II-A Cas9 proteins known to recognize 5′-NGG-3′ PAMs based on the bidentate hydrogen-bonding interactions with guanine nucleotides (Fig. 1C). Because the PAM-recognition mechanism of spCas9 has been elucidated based on its crystal structure (Fig. 1C) (19), BLASTP searches were performed using the spCas9 sequence to identify sequences with distinct conserved PIDs. All retrieved streptococcal Cas9 protein sequences were analyzed by pairwise alignments with the query sequence (spCas9), resulting in identification of other conserved motifs (NQKQ in SmutCas-2, 66.45% identity, and SQSSVR in SoraCas9-2, 66.76% identity). For both pairwise alignments, a distinct divergence was present in the C-terminal PID (≤55% identity). The N-terminal region comprises ~1100 residues and RuvC, REC, and HNH domains (referred to as the “catalytic module” in this study) and shared nearly ~70% identity. We speculated that the PID in the C-terminus was likely acquired by lateral gene transfer. Another BLASTP searches performed using the sequence of the SmutCas-2 catalytic module revealed the highest similarity in the sequence of the catalytic module with that of SmutCas9-1 (93.89% identity), whereas only 43.90% identity was observed for the PID, with distinct NQKQ and DRKR PAM-binding motifs between them (Fig. 2). Using SmutCas-2 PID sequence as query sequence, revealed the highest identity between SmutCas-2 and SmutCas-5 (99.61% identity) as compared with 76.56% identity in their catalytic module (Fig. 2). In summary, these results ruled out full-length gene duplication between SmutCas-2, SmutCas9-1 and SmutCas-5 and strongly indicated that the PID of SmutCas-2 was obtained through domain swapping with SmutCas-5.

**Figure 1.**
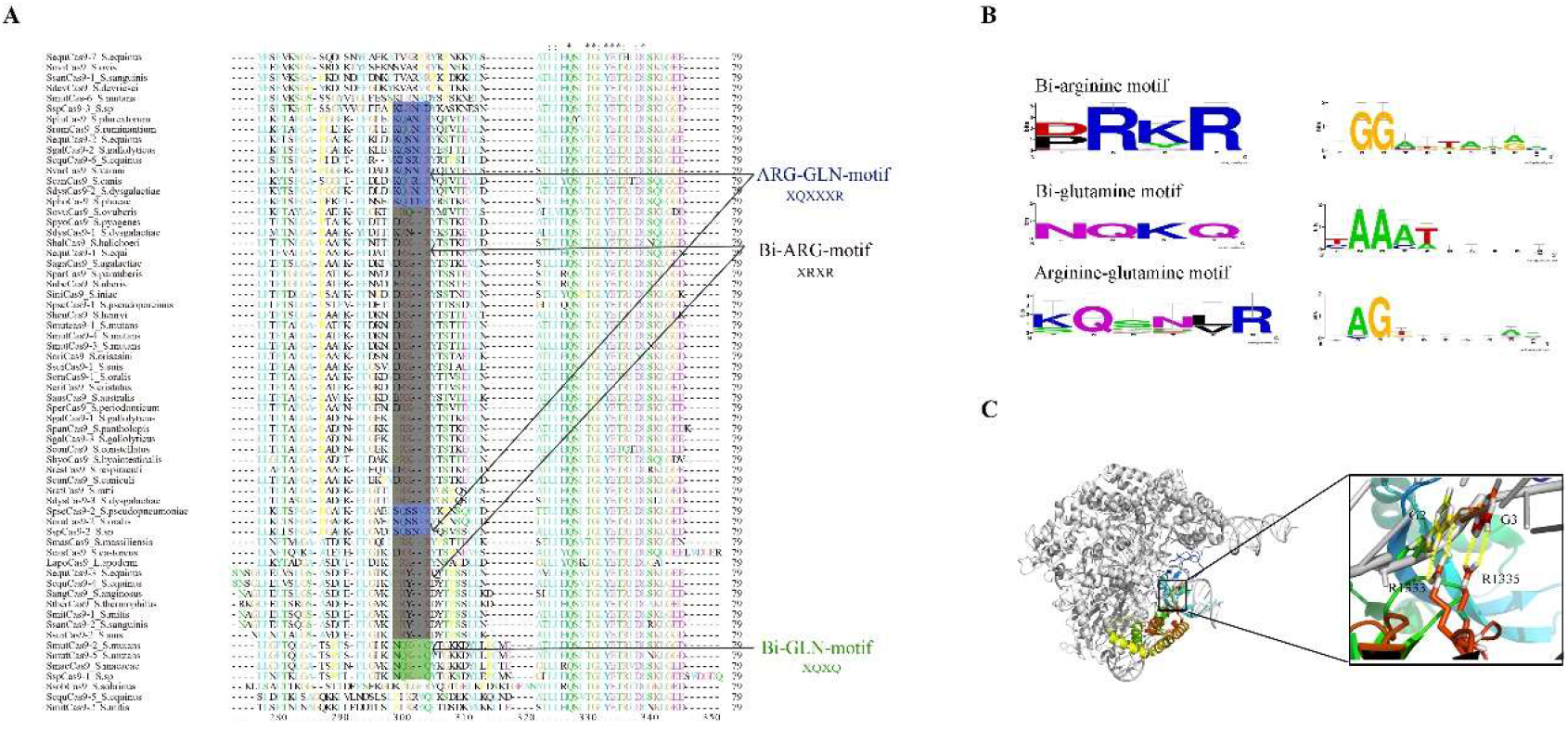
Identification of conserved PID motifs and PAM from natural PAM divergence using bioinformatics. *A*, Multiple sequence alignment of the PAM-binding loops in Cas9 PIDs from the 64 members of genus *Streptococcus*. Multiple sequence alignments were prepared with ClustalX using default colors (orange for the residues G, P, S, and T; red for H, K, and R; blue for F, W, and Y; and green for I, L, M, and V). The bi-arginine, bi-glutamine, and arginine–glutamine motifs are shaded in grey, green, and blue, respectively. *B*, Sequence logo generated online (WebLogo) from inputs of putative PAM sequences found in a *Streptococcus* phage and associated with the bi-arginine, bi-glutamine, and arginine–glutamine motifs. *C*, Domain organization of SpCas9 and a close-up view of PAM-binding sites. A rainbow color map was used for labeling the Cas9 PID. Arginine residues making sequence-specific contacts with the PAM are shown as sticks. Hydrogen-bonding interactions are indicated with dashed lines. PAM, protospacer adjacent motif; PID, PAM-interacting domain; SpCas9, *Streptococcus pyogenes* Cas9.

**Figure 2.**
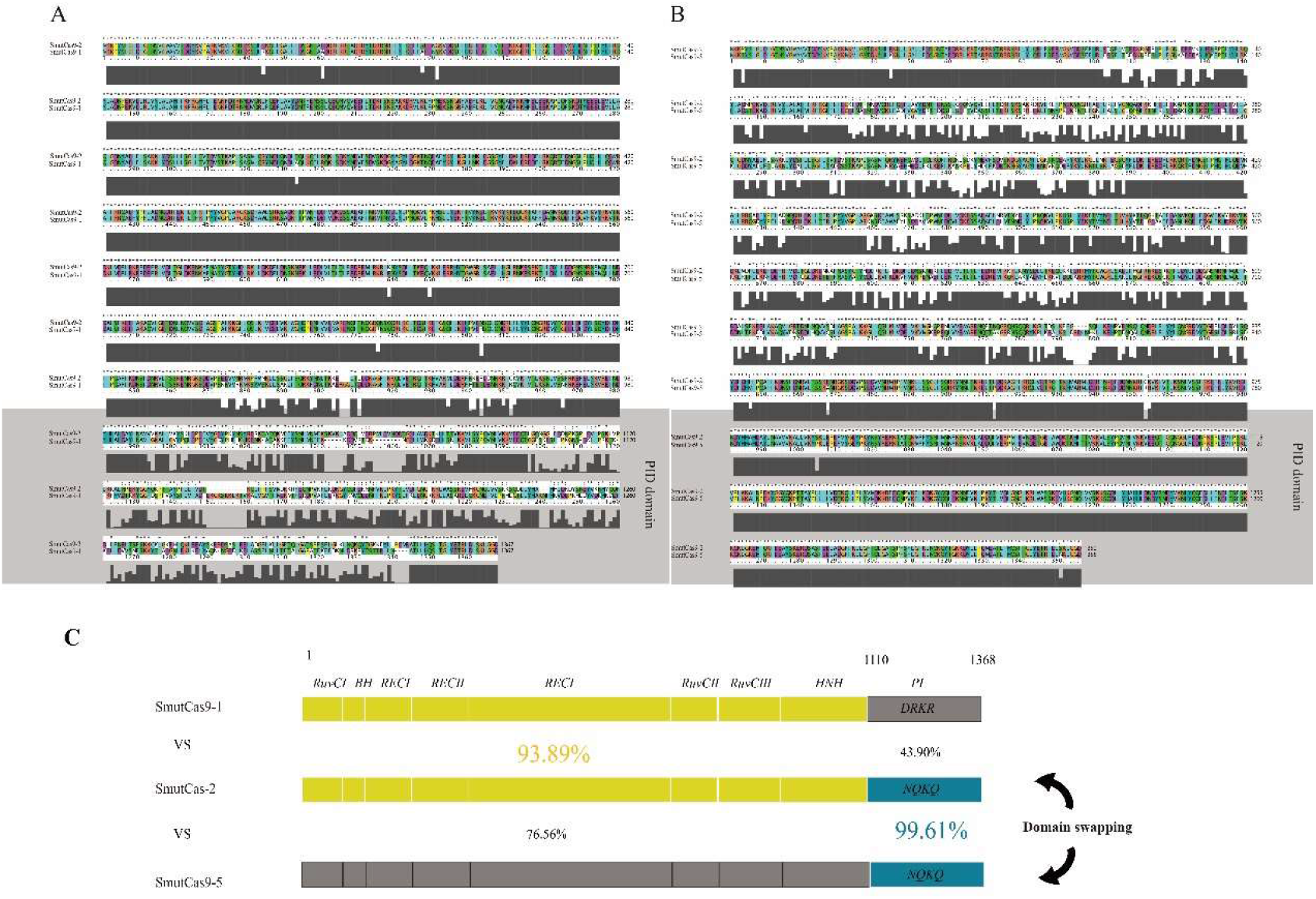
PID exchange between SmutCas9-2 and SmutCas9-5. *A* and *B*, Global pairwise sequence alignment of SmutCas9-2 vs. SmutCas9-1 and SmutCas9-2 vs. SmutCas9-5. Column score is shown in the grey column and indicates the number of correct alignments according to the PAM250 scoring matrix. *C*, Sequence identity for catalytic module and PID domain between SmutCas9-1, SmutCas9-2 andSmutCas9-5. PAM, protospacer adjacent motif; PID, PAM-interacting domain.

To analyze conserved features in the PID, Cas9 orthologues from the genus *Streptococcus* were downloaded from RefSeq, resulting in 64 different Cas9 sequences (Table S1). Three types of conserved motifs were identified from multiple alignment of the PID: a bi-arginine motif (XRKR; X = D or P), a bi-glutamine-motif (NQKQ), and a glutamine–arginine motif (XQXXXR; where X is always an amino acid with a small side chain) (Fig. 1). The sequence features agree with the key roles of arginine and glutamine in DNA binding. Arginine residues are commonly used by DNA-binding proteins to recognize guanines in the major groove, whereas adenine interactions typically involve glutamine residues (20). Additionally, we found that the motifs were not strictly correlated with the species. For example, the NQKQ motif was shared with Cas9s from *Streptococcus macacae* and *Streptococcus mutans*, whereas Cas9s from *S. mutans* harbors both the XRKR and NQKQ motifs.

### Recombinant analysis of Cas9 genes

We then analyzed the sequences of Cas9 genes from *Streptococcus* for possible recombinations. Initial results using an updated version of RDP (21) revealed a propensity for recombination within the *cas9* gene family, as most the sequences were found to have been affected by recombination (Fig. 3). The results showed two features of Cas9s recombination: recombination breakpoints are distributed throughout protein-coding sequences, and breakpoints that fall within domain-coding regions preferentially associate with PIDs (e.g., its midpoint or boundaries). We then analyzed these recombination events by generating and comparing phylogenetic trees from the full-length Cas9 sequence and that of the PID.

**Figure 3.**
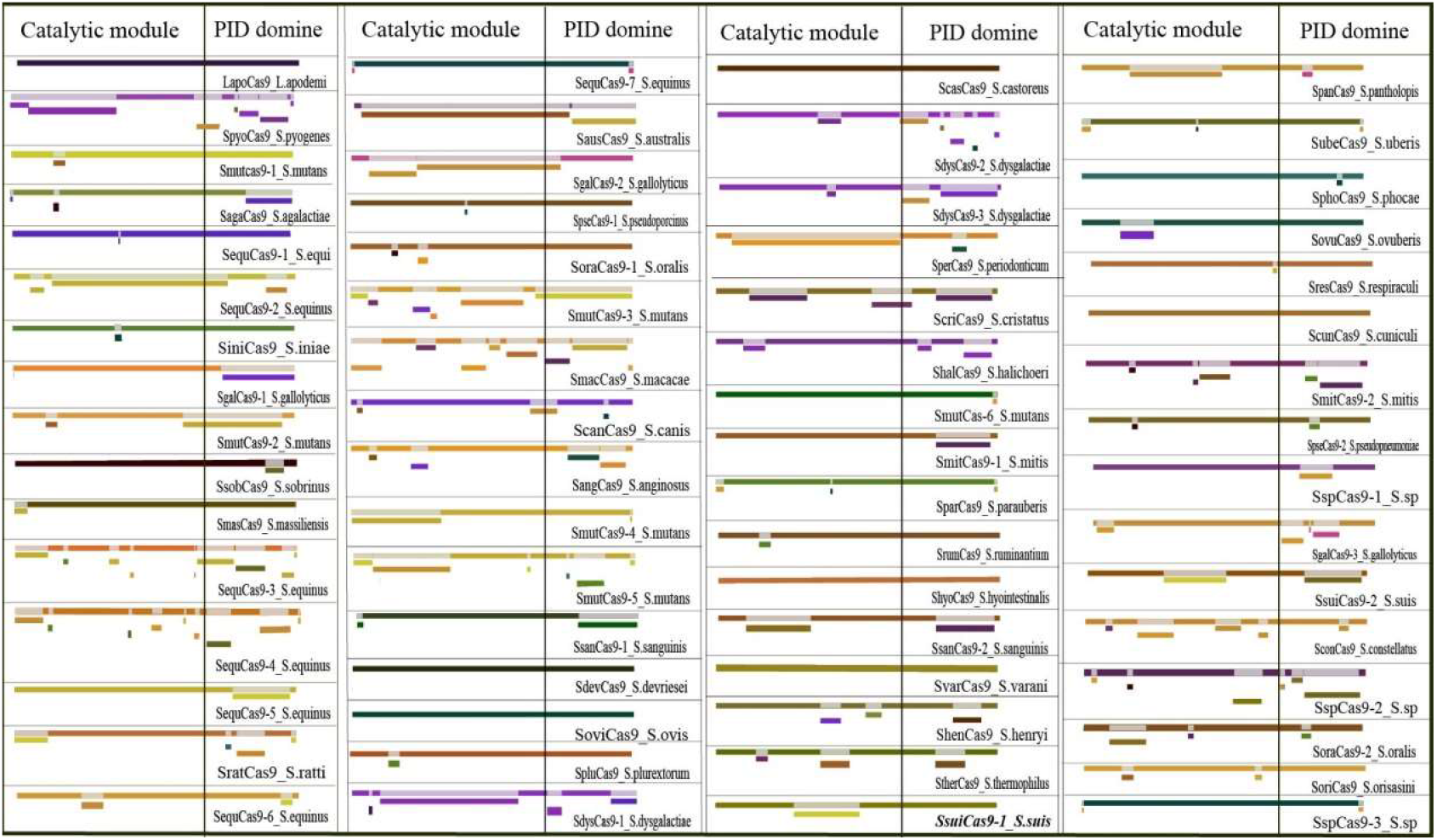
RDP-generated overview of the detected recombination events. The recombination events detected by RDP are indicated below each sequence. The vertical line indicates the borders of the catalytic module and PID. The figure offers an overview of the recombination events in the *cas9* gene dataset. PAM, protospacer adjacent motif; PID, PAM-interacting domain; RDP, recombination detection program.

### Phylogenetic analysis of full-length Cas9 and PID sequences

The aligned Cas9 sequences were analyzed using the MrBayes, and an uprooted consensus tree (Fig. 4) was obtained following Bayesian inference (2,000,000 generations) using a fixed rate and mixed amino acid model. The placements of most branches were supported by high posterior probabilities (in most cases >90%), thereby supporting the reliability of the obtained phylogram.

**Figure 4.**
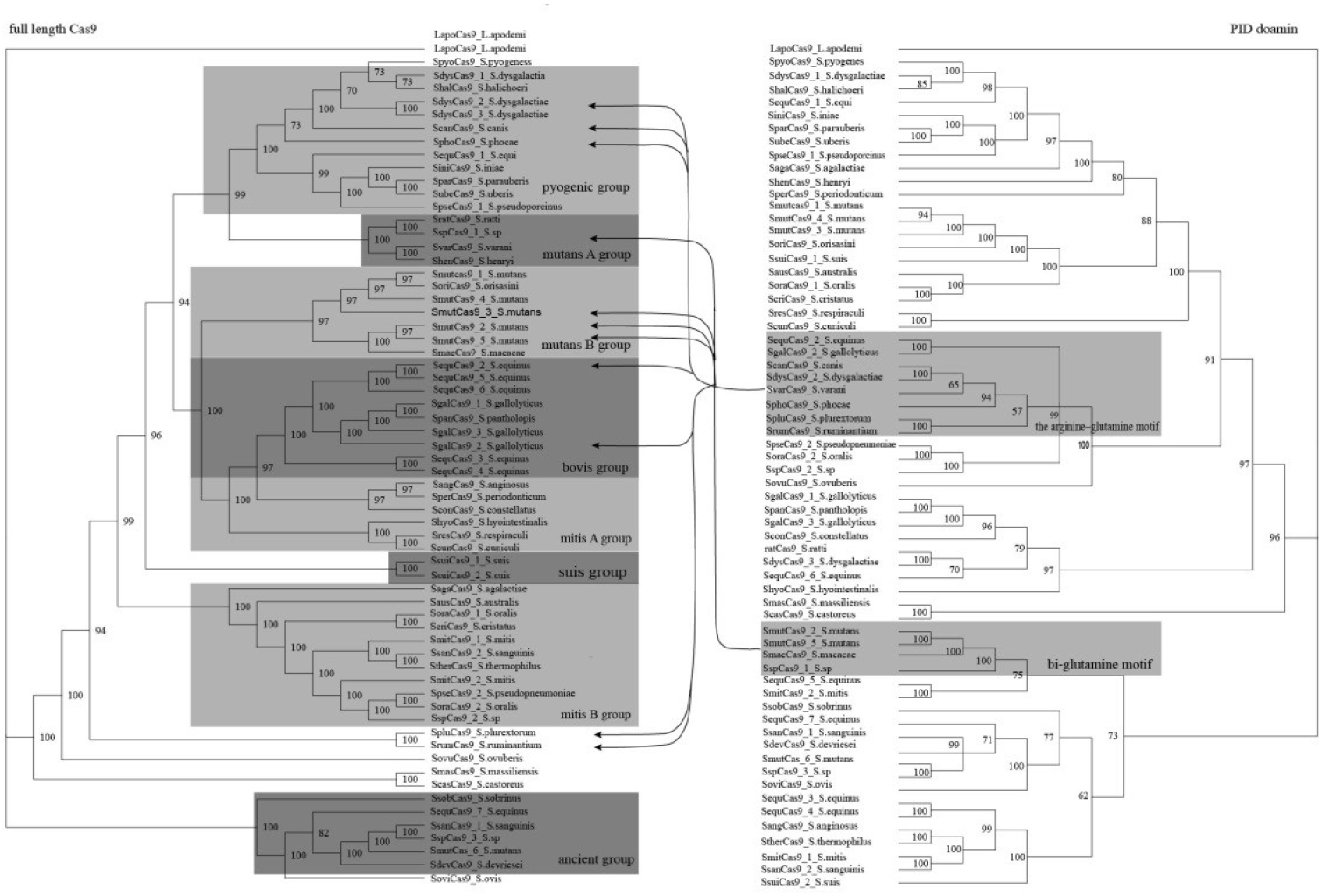
Phylogenic analysis of full-length Cas9 and PID sequences. LapoCas9 from Lactobacillus apodemi was used as outgroup. Boxes indicate the different lineages, and the name of the lineage is displayed in the bottom right corner. Arrows indicate lateral gene transfers of the PID between species. The numbers associated with each node refer to the Bayesian posterior probabilities, shown as percentages. PAM, protospacer adjacent motif; PID, PAM-interacting domain.

The overall topology was similar to that reported for the 16S rRNA coding-region by Kawamura et al. (22). The topology showed six statistically well-supported groups: pyogenic, anginosus, mitis, salivarius, bovis, and mutans (all with posterior probabilities > 95%). Generally, phylogenies inferred either from nucleotide or amino acid sequences did not differ either in major-clade relationships or statistical confidence (i.e., posterior probabilities).

The pyogenic group comprised proteins from *Streptococcus pyogenes, Streptococcus dysgalactiae, Streptococcus halichoeri, Streptococcus canis, Sphocae, Streptococcus iniae, Streptococcus parauberis, Streptococcus uberis*, and *Streptococcus pseudoporcinus*. This large group of proteins also included one protein from *Streptococcus equi*, suggesting lateral transfer of the corresponding gene(s). The mutans group was split into two non-monophyletic clades (mutans A and B). The mutans A group included proteins from *Streptococcus ratti, Streptococcus varani*, and *Streptococcus henryi* and formed a sister cluster with the pyogenic group. The mutans B group included two proteins from *Streptococcus orisasini* and *S. macacae* and five from *S. mutans*. The bovis group comprised five Cas9 proteins from *Streptococcus equinus*, three proteins from *Streptococcus gallolyticus*, and one Cas9 protein from *Streptococcus pantholopis* to form a sister cluster with the mitis group, which was also non-monophyletic taxon. The mitis group appeared to include at least two clusters with proteins from *Streptococcus anginosus, Streptococcus periodonticum Streptococcus constellatus, Streptococcus hyointestinalis, Streptococcus respiraculi, Streptococcus cuniculi, Streptococcus australis, Streptococcus oralis, Streptococcus cristatus, Streptococcus mitis, Streptococcus sanguinis*, and *Streptococcus pseudopneumoniae*. Two *Streptococcus suis* Cas9 proteins formed the small suis group, and one *Streptococcus agalactiae* Cas9 (pyogenic group) protein and one *Streptococcus thermophilus* Cas9 protein (salivarius group) was arranged among the large mitis group. Six Cas9 proteins belonging to four different groups were located outside of the six identified groups; therefore, we speculated that these formed a basal group for the genus and named it the ancient group. A possible scenario explaining the observed mosaic distribution in this group could be that a *cas9* gene might have arisen early and undergone a lateral transfer. This cluster might represent the most ancient group of Cas9s in the *Streptococcus* genus.

The resulting tree topology from the PID sequence was clearly dominated by the conserved PAM-interacting motifs rather than species phylogeny (Fig. 4). Although this tree did not form statistically well-supported groups, expect for one pyogenic group and one mutans group, the existence of the conserved motif determining the substrate specificity was well reflected by this phylogeny. According to multiple alignment results, *Streptococcus* Cas9 PIDs might be further classified into three types with bi-arginine, bi-glutamine, and arginine–glutamine motifs. A closer look at the location of these PID domains revealed that all of three typical motifs formed groups with well supported. Additionally, the different distribution patterns of the trees generated from the full-length and PID sequences could be explained by domain-exchange events between Cas9 sequences. For an example, the arginine–glutamine motif group was formed by the PID sequences of SseqCas9-2, SgalCas9-2, ScanCas9, SdysCas9, SvarCas9, SphoCas9, SpluCas9, and SrumCas9. Phylogenetic analysis of full-length Cas9 indicated that these Cas9 proteins formed distinct groups (ScanCas9, SdysCas9, and SphoCas9 in pyogenic; SvarCas9 in mutans group; SseqCas9-2 and SglaCas9-2 in bovis; and SpluCas9 and SrumCas9 in suis). Both genera and conserved-motif groups formed a stable cluster (≥95% posterior probabilities). These findings strongly supported a naturally occurring PID domain-shuffling process among *cas9* genes.

### PAM specificities with conserved PAM-interacting motifs

To identify the functional PAMs for Cas9s harboring bi-arginine, bi-glutamine, or arginine– glutamine motifs, potential protospacers matching spacer sequences were searched as described in methods. We aligned the identified 10-nt sequences located directly downstream of the protospacer sequence and delineated the most common nucleotides that could represent PAM sequences. The identified PAM specificities agreed well with well-characterized amino acid–base recognition commonly involving arginine and glutamine residues. The potential PAMs of the bi-arginine, bi-glutamine, and arginine–glutamine motifs were NGG, NAA, and NAG, respectively (Fig. 2B). Notably, the positions of the arginines and glutamines in the bi-arginine and bi-glutamine motifs are well aligned; however, previous attempts to change the PAM specificity of SpCas9 to NAA by introducing arginine-to-glutamine substitutions were unsuccessful (19). These results suggest that arginine-to-glutamine substitutions within a given motif are necessary but insufficient to change the specificity from guanines to adenines. Multiple rounds of somatic mutation are likely needed to realize a change in PAM specificity. Unlike random mutagenesis, when facing novel phages, PID exchange represents a rapidly evolving strategy that is likely a consequence of an “arms race” against rapidly evolving viruses.

### The distribution of PID-swapping events across the Cas9 family

To further analyze PID-swapping distributions among bacterial Cas9s, we used Cas9 protein sequences as a query sequence to search the RefSeq non-redundant protein database to discover orthologues with shared divergent PIDs resulting from domain swapping. Table 1 shows PID-swapping events according to genus (containing >10 members for each sequence) and occurring in >50% of the genera. Of the 582 Cas9s identified, 53 demonstrated PID swapping (hit rate: 1–10), with the number of PID-swapping events per genus varying from zero to fourteen (Table 1). Additionally, 11 Cas9s demonstrated PID swapping between genera. These results clearly demonstrated that PID swapping broadly occurred throughout the Cas9 family.

**Table 1.**
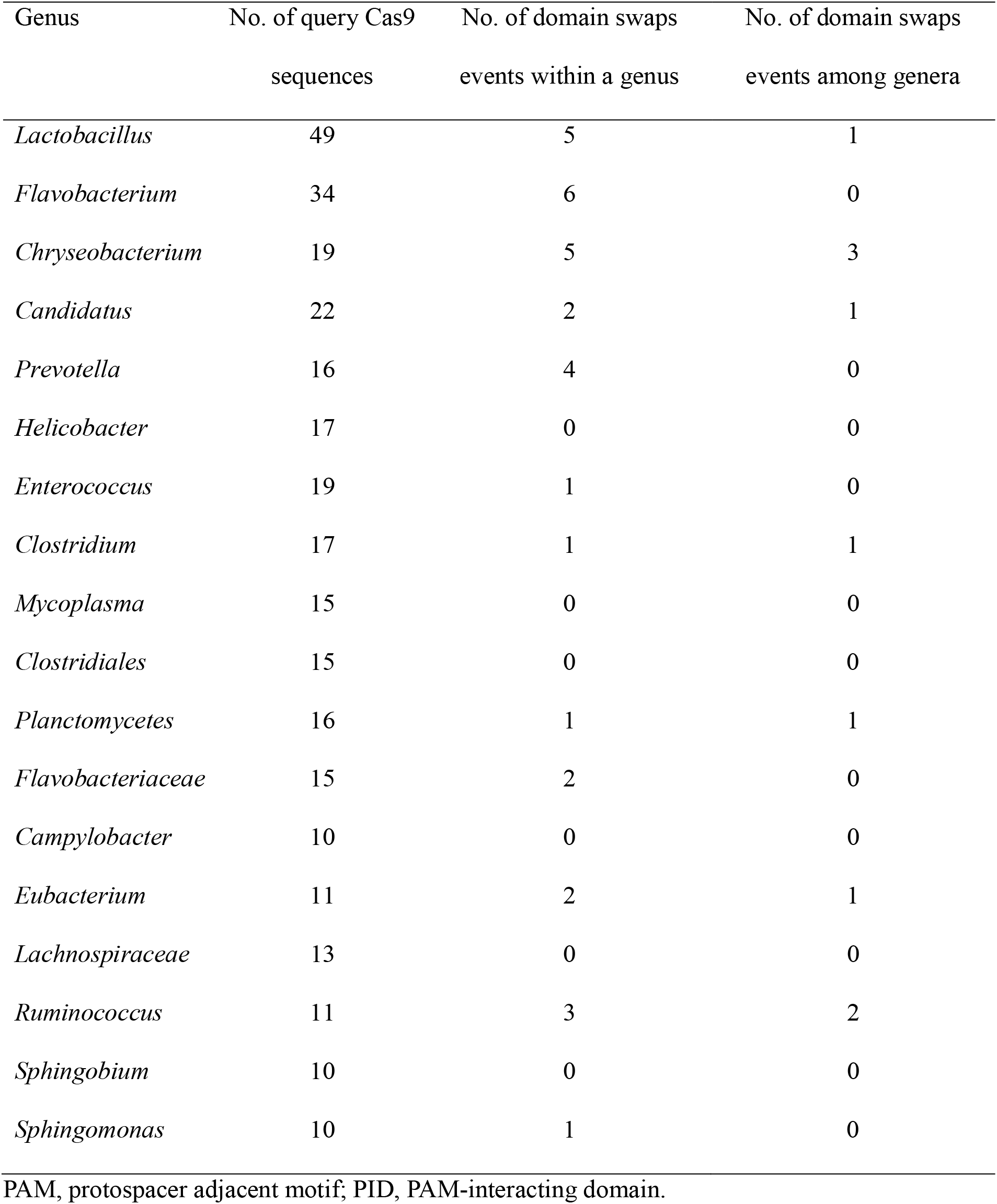
The distribution of PID-swapping events for selected Cas9 genera.

| Genus | No. of query Cas9<br>sequences | No. of domain swaps<br>events within a genus | No. of domain swaps<br>events among genera |
| --- | --- | --- | --- |
| *Lactobacillus* | 49 | 5 | 1 |
| *Flavobacterium* | 34 | 6 | 0 |
| *Chryseobacterium* | 19 | 5 | 3 |
| *Candidatus* | 22 | 2 | 1 |
| *Prevotella* | 16 | 4 | 0 |
| *Helicobacter* | 17 | 0 | 0 |
| *Enterococcus* | 19 | 1 | 0 |
| *Clostridium* | 17 | 1 | 1 |
| *Mycoplasma* | 15 | 0 | 0 |
| *Clostridiales* | 15 | 0 | 0 |
| *Planctomycetes* | 16 | 1 | 1 |
| *Flavobacteriaceae* | 15 | 2 | 0 |
| *Campylobacter* | 10 | 0 | 0 |
| *Eubacterium* | 11 | 2 | 1 |
| *Lachnospiraceae* | 13 | 0 | 0 |
| *Ruminococcus* | 11 | 3 | 2 |
| *Sphingobium* | 10 | 0 | 0 |
| *Sphingomonas* | 10 | 1 | 0 |

PAM, protospacer adjacent motif; PID, PAM-interacting domain.

### PID domain swapping of Cas9s from Staphylococcus

As previous studies has reported that swapping PID domains between Cas9 proteins of Streptococcus genus generated chimeras with exchanged PAM specificities (23,24). Here, we use SaCas9 protein from *Staphylococcus aureus* as the example to illustrate the streamline to develop a variant with novel PAM by PID domain swapping. The SaCas9 is a compact Cas9 ortholog suitable for viral delivery for biomedical applications, and SaCas9 displays a comparable activity to SpCas9 in mammalian cells (9). The wild-type SaCas9 requires an NNGRRN (R = A or G) PAM with a preference for a T base at the 6th position of the PAM (9). Firstly, using the protein sequence of saCas9 as the query sequence, a BLASTP search revealed one homologous Cas9 (NCBI accession number WP_039643679.1) from *Staphylococcus hyicus*, termed as SHyCas9 in this study, with a divergent PID domain sequence. Secondly, PAM bioinformatic analysis as described in methods was carried out to uncover a novel NNAAAA PAM specificity of *Staphylococcus hyicus* Cas9 (Fig. 5B). Finally, a domain swapped chimera SaHyCas9 was created using the sub-cloning gene fragment of the PID domain of SHyCas9 replacing that of saCas9 (Fig. 5A). We verified the PAM specificity of SaHyCas9 in human cells by cotransfecting human embryonic kidney–293T (HEK293T) cells with a plasmid expressing these variants along with sgRNAs and a plasmid containing target sequence and PAM. After 72 hours post-transfection, gene modification rates as detected by the T7E1 assay. The result that demonstrated SaHyCas9 show the remarkable activity to target sequence with NNAAAA (CTAAAA) PAM comparable with that of SaCas9 with NNGRRT (CTGGAT) PAM sequence (Fig. 5C). These results verify that SaHyCas9 can serve as a potential alternative to SaCas9 for DNA editing in mammalian cells and demonstrated that domain swapping should be an appealing platform to discover novel Cas9 PAM specificities.

**Figure 5.**
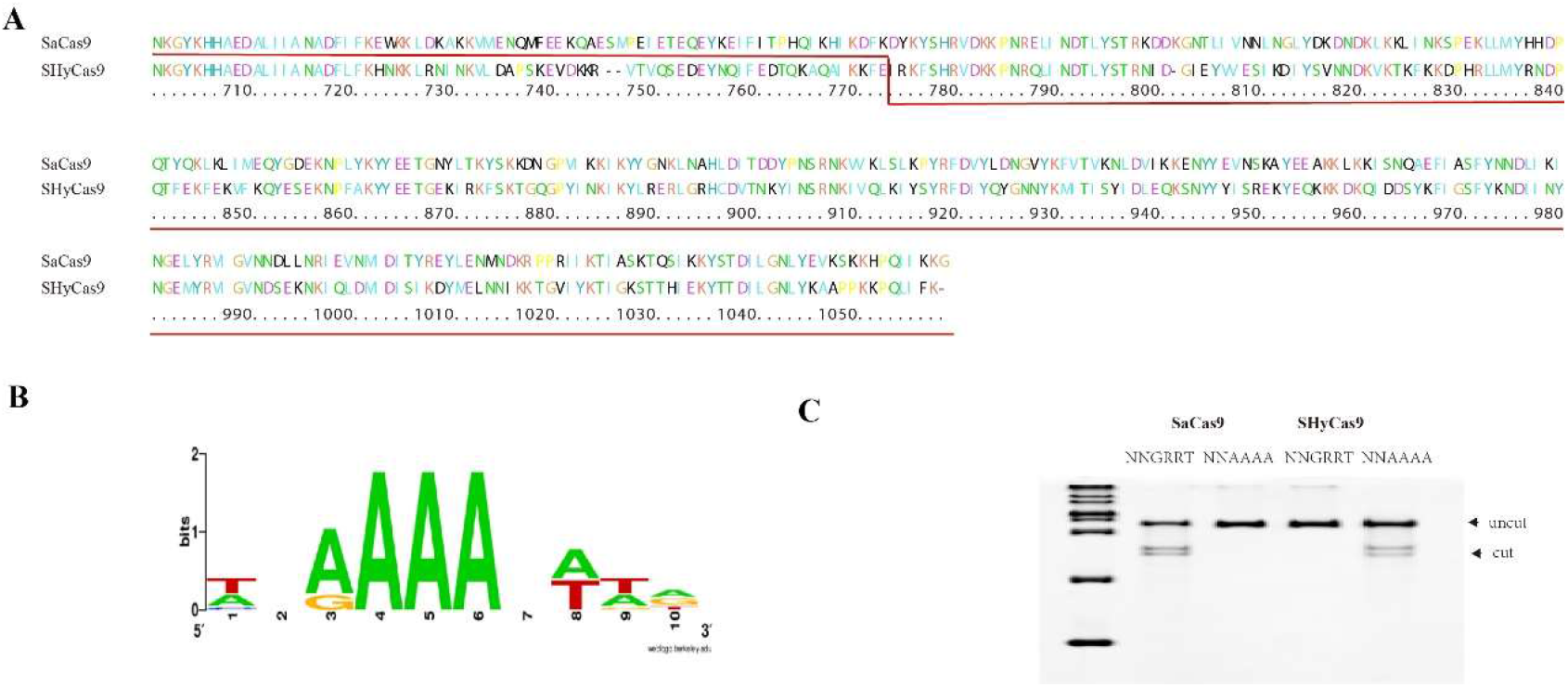
PID domain swapping of Cas9s from Staphylococcus. A sequence alignment of SaCas9 and SaHyCas9. *A*, PID domain pairwise alignment between SaCas9 and SHyCas9. The step in the underlining red line marks the joining of SaCas9 and SHyCas9 to construct a SaHyCas9 hybrid. *B*, Identification of PAM specificity of SHyCas9 through bioinformatics. A sequence logo generated online (WebLogo) that was input with putative PAM sequences found in Staphylococcus plasmids and associated with close SHyCcas9 homologs. *C*, T7E1 analysis of indels produced at plasmid in cellula with indicated PAM. Arrows distinguish banding of the cleaved products from uncleaved substrate (top band). PAM, protospacer adjacent motif; PID, PAM-interacting domain.

## Discussion

Earlier studies have detected CRISPR-based immune systems are essentially modular and appear to exchange readily between closely related organisms (25–27). In this study, we demonstrated that the PID domain exchanging profusely occurs in Cas9 family and play an important role in fast altering its PAM specificity. Recombination analysis revealed that the recombination events at the Cas9 was not limited to the PID domain region. Actually, the recombination events distributed throughout protein-coding sequences of Cas9 sequence (Fig. 3). Phylogenetic analysis of the full-length Cas9 sequence also discovered an ancient Cas9 group spanning over several main group of Streptococcus resulting from lateral gene transfer (Fig. 4). Previous phylogenomic studies had revealed an extensive horizontal transfer of CRISPR locus among bacteria (22). In conclusion, for CRISPR system, exchanging of gene fragments at functional domain, complete gene and locus level should occur frequently and may play a key role in race against rapidly evolving viruses.

DNA cleavage by SpCas9, the most widely used Cas9, is dependent on the presence of a 5’-NGG-3’ PAM in the target DNA, thereby restricting the choice of targetable sequences. To allow SpCas9 access to more of the genome, researchers have performed profound protein engineering in an attempt to generate wider ranges of PAM preference. Directed evolution has been successfully utilized to generate mutants Cas9 with relaxed PAM requirements. However, the prerequisite for sensitive and efficient screening method makes such processes time consuming, thereby limiting access to a broader spectrum of substitutions and hindering wider utilization. Conversely, structure-guided engineering allows mutation of fewer residues but at a greater depth by using structural information as a rational guide; however, there have been few successful reports on rational designing of the Cas9 PAM. This could be the result of an incomplete understanding of the underlying mechanisms required for PAM recognition. Previous attempts to change SpCas9 PAM specificity to an NAA PAM by simply introducing arginine-to-glutamine substitutions at positions 1333 and 1335 were unsuccessful. A plausible explanation is that the Gln1335 side chain, being 1.5-Å shorter than arginine, is unable to reach the adenine base in the major groove without further structural rearrangements. In the present study, sequence alignment revealed the presence of a PCME sequence proximal to the XQXQ motif in the PAM-interacting loop. The distal side chain of glutamine can adopt the same position in major grooves of DNA as arginine due to the increased flexibility introduced by the elongated loop. The arginine–glutamine (XQXXXR) motif includes amino acids with small side chains inserted between the glutamine and arginine, which likely increases the flexibility of the binding loop. These findings demonstrate that in addition to promoting sequence-specific interactions with the PAM, introduction of specific amino acids improved loop flexibility. Both substitutions and insertions-deletions should be explored during rational design of novel PAM-recognition sequences.

The best protein engineering strategies are likely to be those that most closely mimic natural ones. Considering that the creation of a novel chimeric variant with novel PAM specificity, just need a bioinformatics analysis and one step cloning. Domain swapping should be an appealing approach to tuning the Cas9 PAM specificity, given that a plenty diverse exchangeable PID domain existing in natural as revealed by data analysis protein sequence database. In our study, using saCas9 as example, we created a functional chimeric Cas9 with a novel NNAAAA PAM specificity in one week. This evolved mutant also suffers from long PAM of NNAAAA. However this was the first reported saCas9 orthologue with adenine-rich PAM recognition. It will be interesting should be a good starting point for a short protospacer adjacent motif with adenine preference by protein engineering. Moreover, this domain swapping approach could be easily adapted for PAM expanding of other CRISPR–Cas9 nucleases

In summary, studying evolutionary mechanisms could offers insight into how to use the adaptability of proteins and to build new variants. The results of the present study demonstrate that domain swapping in Cas9 should be a powerful approach for extended PAM-binding capabilities. Furthermore, a shuffled PAM-specific PID domain based on a well-characterized scaffold could provide a good starting point for further PAM engineering. Cas9 orthologues are abundant, with >4000 Cas9 sequences currently available. Exploring diversity of PIDs in these Cas9 orthologues might allow the generation of variants with novel PAM specificities and expand the repertoire of Cas9s available for genome-targeting applications.

## Experimental procedures

### Protein sequence analysis

To detect diversely Cas9 orthologues, we used well-characterized SpCas9 as the query sequence to perform BLASTP searches with the default settings (https://blast.ncbi.nlm.nih.gov/Blast.cgi?PAGE=Proteins). Protein sequences containing motifs different from the typical XRXR motifs in SpCas9 were collected for domain-swapping analysis. For multiple alignments and phylogenetic analyses, all Cas9 sequences from streptococcal bacteria were retrieved from the NCBI reference database (https://www.ncbi.nlm.nih.gov/refseq/). We manually removed sequences with incomplete open reading frames, and redundant sequences (90% identity) according to the sequence demarcation tool (http://web.cbio.uct.ac.za/~brejnev/) among all streptococcus Cas9 sequences were removed (21). This procedure identified 63 Cas9 sequences, with the accession numbers shown in Table S1.

### Recombination analysis

The nucleotide sequence from the coding regions of selected Cas9 variants was download from the NCBI nucleotide database (https://www.ncbi.nlm.nih.gov/nucleotide), aligned using ClustalX, and manually corrected (28). To identify recombination events, we used a modified version of the recombination detection program (RDP) that implements additional methods for detecting recombinations (29). The BOOTSCAN algorithm (30) was used to check the initial RDP results. The parameters used for RDP analysis were as follows: no multiple-comparison correction, a highest acceptable *p*-value of 0.001, and a reference sequence selection using internal references.

### Phylogenetic analysis

Bayesian inference analysis was performed on the matrix using MrBayes (v.3.7) (31). For DNA phylogenetic analysis, we selected a DNA-substitution model from a comparison of 32 models using the Akaike information criterion as implemented in MrModeltest (v.2.3) (32). For protein phylogenetic analysis, priors included a mixed amino acid model allowing for optimization of the model during analysis (31). The Cas9 sequence of Lactobacillus apodemi served as an outgroup in both DNA and protein phylogenetic analyses. MrBayes analyses were run with the following parameters: mcmcngen = 2,000,000, samplefreq = 1000, nchains = 4, startingtree = random and sumt burnin = 250. Split frequencies were checked to ensure convergence. The resulting Bayesian trees were visualized using the program Figtree (http://tree.bio.ed.ac.uk/software/figtree/).

### Searching for PAM motifs

Spacer sequences of the selected bacterial species were extracted from the CRISPR database (http://crispr.u-psud.fr/crispr/) and used to find protospacer candidates using BLAST (http://blast.ncbi.nih.gov/Blast). Protospacer candidates were defined as those containing a phage DNA sequence with ≥90% similarity to the crRNA spacer. A logo plot (http://weblogo.berkeley.edu/) showing the most abundant nucleotides was created, and PAM sequences were predicted. The spacer sequences were then used to select cognate protospacer sequences, as described.

### Analysis of PID exchange within the entire Cas9 family

All Cas9 protein sequences were retrieved from the UniProtKB database (https://www.uniprot.org/) using “Cas9” as the search term. Sequences <800 amino acids were discarded as truncated Cas9 variants. To remove redundancy, the results in sequence clusters with 90% identity were collected. The selected sequences were clustered according to genus. As expected, genera with a large number of members often showed a higher level of domain-swapping activity that those fewer members; therefore, those with <10 members were discarded and not used for further domain-swapping analyses. 582 representative sequences (provided in Data S1) were used as queries to perform BLASTP searches. The pairwise aligned sequences with homologous catalytic module (>80% identity) and a divergent PID sequence (<60% identity) were defined as PID-swapping events.

### Plasmids and oligonucleotides

Sequences of plasmids and oligonucleotides used in this study can be found in Data S2. The gene fragment encoding the PAM-interaction domain of S. hyicus, synthesized by Generay (Shanghai, China), was inserted into a vector PX601-containing wild-type SaCas9 (Addgene plasmid #61591) and replacing that domain of SaCas9. The resulting hybrid SauHyi Cas9 construct was sequence-verified by a next-generation sequencing service (Generay Biotech, China). The 646 bp target DNA containing target site sequence, the PAM sequence and green fluorescent protein encoding sequence was synthesized and inserted in XbaI/SphI-digested PLV vector. Plasmid containing NNGRRT (CTGGAT) and NNAAAA (CTAAAA) termed as pLV-GRRT and pLV-AAAA, respectively. Transfection efficiencies as measured by the expression of a green fluorescent protein. For verification of PAM sequence by cleavage of plasmid DNA in cellular, 50 ng crRNA and nuclease expression plasmid and 50 ng PAM sequence containing plasmids pLV-GRRT or pLV-AAAA.

### Verification of PAM sequence by cleavage of plasmid DNA in cellula

HEK293T cells were cultured in high-glucose DMEM complete media (Dulbecco’s modified Eagle’s medium (DMEM)), 4.5 g/L glucose, 0.045 unit/mL of penicillin, 0.045 g/mL streptomycin, and 10% FBS (Sangon Biotech, China) at 37°C, 100% humidity, and 5% CO2. One day before transfection, ~2 × 105 HEK293T cells in 1 mL of high-glucose DMEM complete media were seeded into each well of 12-well plastic plates (Corning). Shortly before transfection, the medium was replaced with fresh DMEM complete media. The transfection experiments were performed by using Hieff TransTM Liposomal Transfection Reagent (Yeasen, China) by following the manufacturer’s protocol. Each transfection experiment was independently repeated.

HEK293T cells were transfected with plasmids, as described above. Cells were incubated at 37°C for 60 h after transfection before DNA extraction. Cellula DNA was extracted using the TIANamp Genomic DNA Kit (TIANGEN, China) following the manufacturer’s protocol. The plasmid DNA region flanking the CRISPR target site was PCR amplified (primers listed in Data S2). 500 ng total of the purified PCR products were mixed with 1 μl 10× NEBuffer 2 (New England Biolabs Inc., Beverly, MA, USA) and ultrapure water to a final volume of 9.75 μl, and subjected to a re-annealing process to enable heteroduplex formation: 95°C for 2 min, 95°C to 85°C ramping at −2°C/s, 85°C to 25°C at −0.1°C/s, and 16°C hold for 1 min. After re-annealing, products were treated with 0.25 μl T7 Endonuclease I enzyme, incubated for 20 min at 37°C, and analyzed on 10% Giass gei® UREA-TBE PAGE (WSHT). Gels were stained with 0.5μg/mL Ethidium Bromide (Sangon Biotech, china) for 20 min and imaged with a Gel image system (Tanonc, china).

### Statistical analysis

For BLAST and recombination analysis, a P-value <0.001 was considered to be the cut-off criterion for significance. For phylogenetic analysis, the Bayesian posterior probabilities were used to indicate branch supports. Posterior probabilities > 90% indicated a good support.

## Supporting information

sportting information

## Data availability

All data needed to evaluate the conclusions in the paper are present in the paper and the Supporting information.

## Supporting information

This article contains supporting information.

## Funding and additional information

This work was supported by the Natural Science Foundation of Zhejiang Province No. LY17C050001, and the National Natural Science Foundation of China Nos. 81730108 and 8197363.

## Abbreviations

crRNA: CRISPR RNA
DMEM: Dulbecco’s modified Eagle’s medium
PAM: protospacer adjacent motif
PID: PAM-interacting domain
RDP: recombination detection program
SpCas9: *Streptococcus pyogenes* Cas9
SpRY: SpCas9 variant
SaCas9: *Staphylococcus aureus* Cas9
SHyCas9: *Staphylococcus hyicus* Cas9.

