## Supplementary material for "PAM-interacting domain swapping is extensively utilized in nature to evolve CRISPR-Cas9 nucleases with altered PAM specificities": sportting information

*From the [1]College of Pharmacy, School of Medicine, Hangzhou Normal University, Hangzhou, Zhejiang 311121, China; [2]Key Laboratory of Elemene Class Anti-Cancer Chinese Medicines; Engineering Laboratory of Development and Application of Traditional Chinese Medicines; Collaborative Innovation Center of Traditional Chinese Medicines of Zhejiang Province, Hangzhou Normal University, Hangzhou, Zhejiang 311121, China; [3]College of Basic Medicine, School of Medicine, Hangzhou Normal University, Hangzhou, Zhejiang 311121, China*

**This PDF file includes:**

Table S1

Data S2

**Other Supplementary information for this manuscript includes the following:**

Data S1

**Table S1. The sixty-four Cas9s analyzed in this study.**

| Name | Host | NCBI Reference SequenceAccession no. | Binding motif in PID domain |
|---|---|---|---|
| LapoCas9 | Lactobacillus apodemi | WP_025088082.1 | ND |
| SpyoCas9 | Streptococcus pyogenes | WP_002989955.1 | DRKR |
| SmutCas9-1 | Streptococcus mutans | WP_002277364.1 | DRKR |
| SagaCas9 | Streptococcus agalactiae_ | WP_001040078.1 | DRKR |
| SequCas9-1 | Streptococcus equi | WP_012515931.1 | DRKR |
| SequCas9-2 | Streptococcus equinus | WP_004232481.1 | KQSNLR |
| SiniCas9 | Streptococcus iniae | WP_003099269.1 | DRKR |
| SgalCas9-1 | Streptococcus gallolyticus | WP_012962174.1 | PRKR |
| SmutCas9-2 | Streptococcus mutans | WP_002308623.1 | NQKQ |
| SsobCas9 | Streptococcus sobrinus | WP_019783835.1 | ND |
| SmasCas9 | Streptococcus massiliensis | WP_018372492.1 | PRKR |
| SequCas9-3 | Streptococcus equinus | WP_115255320.1 | PRYR |
| SequCas9-4 | Streptococcus equinus | WP_074965192.1 | PRYR |
| SequCas9-5 | Streptococcus equinus | WP_074639375.1 | ND |
| SratCas9 | Streptococcus ratti | WP_003088697.1 | PRKR |
| SequCas9-6 | Streptococcus equinus | WP_074481527.1 | KISRIR |
| SequCas9-7 | Streptococcus equinus | WP_074483649.1 | ND |
| SausCas9 | Streptococcus australis | WP_125389260.1 | ERKR |
| SgalCas9-2 | Streptococcus gallolyticus | WP_077497300.1 | KQSNLR |
| SpseCas9-1 | Streptococcus pseudoporcinus | WP_007896501.1 | DRKR |
| SoraCas9-1 | Streptococcus oralis | WP_084868974.1 | DRKR |
| SmutCas9-3 | Streptococcus mutans | WP_002304487.1 | DRKR |
| SmacCas9 | Streptococcus macacae | WP_003079701.1 | NQKQ |
| ScanCas9 | Streptococcus canis | WP_003043819.1 | KQGRLR |
| SangCas9 | Streptococcus anginosus | WP_003031511.1 | PRYR |
| SmutCas9-4 | Streptococcus mutans | WP_002264920.1 | DRKR |
| SmutCas9-5 | Streptococcus mutans | WP_002277050.1 | NQKQ |
| SsanCas9-1 | Streptococcus sanguinis | WP_002933589.1 | ND |
| SdevCas9 | Streptococcus devriesei | WP_027976660.1 | ND |
| SoviCas9 | Streptococcus ovis | WP_018377631.1 | ND |
| SpluCas9 | Streptococcus plurextorum | WP_027972728.1 | KQANLR |
| SdysCas9-1 | Streptococcus dysgalactiae | WP_126426708.1 | KRNR |
| ScasCas9 | Streptococcus castoreus | WP_027970237.1 | DRRR |
| SdysCas9-2 | Streptococcus dysgalactiae | WP_155782941.1 | KQGNLR |
| SdysCas9-3 | Streptococcus dysgalactiae | WP_143927572.1 | PRKR |
| SperCas9 | Streptococcus periodonticum | WP_174720700.1 | DRKR |
| ScriCas9 | Streptococcus cristatus | WP_149518150.1 | DRKR |
| ShalCas9 | Streptococcus halichoeri | WP_159583349.1 | DRKR |
| SmutCas-6 | Streptococcus mutans | WP_168745320.1 | PRYR |
| SmitCas9-1 | Streptococcus mitis | WP_101785820.1 | PRYR |
| SparCas9 | Streptococcus parauberis | WP_103344984.1 | DRKR |
| SrumCas9 | Streptococcus ruminantium | WP_156010052.1 | KQGNLR |
| ShyoCas9 | Streptococcus hyointestinalis | WP_115268905.1 | PRKR |
| SsanCas9-2 | Streptococcus sanguinis | WP_176798877.1 | PRYR |
| SvarCas9 | Streptococcus varani | WP_093650272.1 | KQSNLR |
| ShenCas9 | Streptococcus henryi | WP_074484960.1 | DRKR |
| StherCas9 | Streptococcus thermophilus | WP_024703962.1 | PRYR |
| SsuiCas9-1 | Streptococcus suis_ | WP_160864239.1 | DRKR |
| SpanCas9 | Streptococcus pantholopis | WP_067062573.1 | PRKR |
| SubeCas9 | Streptococcus uberis | WP_111673267.1 | ERKR |
| SphoCas9 | Streptococcus phocae | WP_054279288.1 | KQDDVR |
| SovuCas9 | Streptococcus ovuberis | WP_168548033.1 | PRQR |
| SresCas9 | Streptococcus respiraculi | WP_119877030.1 | DRKR |
| ScunCas9 | Streptococcus cuniculi | WP_075103982.1 | DRKR |
| SmitCas9-2 | Streptococcus mitis | WP_061590506.1 | ND |
| SpseCas9-2 | Streptococcus pseudopneumoniae | WP_049549711.1 | SQSSVR |
| SspCas9-1 | Streptococcus sp | WP_162011508.1 | NQKQ |
| SgalCas9-3 | Streptococcus gallolyticus | WP_058692367.1 | PRKR |
| SsuiCas9-2 | Streptococcus suis_ | WP_074391585.1 | PRYR |
| SconCas9 | Streptococcus constellatus | WP_048800889.1 | PRKR |
| SspCas9-2 | Streptococcus sp | WP_067192512.1 | SQSNVR |
| SoraCas9-2 | Streptococcus oralis | WP_061587801.1 | SQSSVR |
| SoriCas9 | Streptococcus orisasini | WP_057491067.1 | DRKR |
| SspCas9-3 | Streptococcus sp | WP_070669353.1 | KLRNVD |

Cas9 names should include an organism (1 letter genus abbreviation and 3 letter species abbreviation) and gene designation. A given species may contain one or several protein sequences; within the species, the sequences are numbered using Arabic numerals.
ND, not determined.

**Data S2.** Sequences of plasmids and oligonucleotides used in this study.

**Plasmids used in this study**

| **Name** | **Addgene ID** | **Description** |
|---|---|---|
| PX601 | 61591 | CMV-NLS-SaCas9-NLS-3xHA;U6-BsaI-SagRNA |
| PX601-Sacas9-hpfh1 | ---- | CMV-NLS-SaCas9-NLS-3xHA;U6-hpfh1- SagRNA |
| PX601-SaHycas9-hpfh1 | ---- | CMV-NLS- SaHycas9-NLS-3xHA;U6-hpfh1- AagRNA |
| pLV-puro | 125839 | CMV |
| pLV-NNGRRT | ---- | CMV-hpfh1-NNGRRT-GFP |
| pLV-NNAAA | ---- | CMV-hpfh1-NNAAAA-GFP |

PX601: CMV-NLS-SaCas9-NLS-3xHA;U6-BsaI-sgRNA

CMV promoters colored in green, human codon optimized S. aureus Cas9 colored in blue, NLS underlined, U6 promoters colored in yellow, BsaI sites underlined, SagRNA colored in purple.

ggtgatgcggttttggcagtacatcaatgggcgtggatagcggtttgactcacggggatttccaagtctccaccccattgacgtcaatgggag
tttgttttggcaccaaaatcaacgggactttccaaaatgtcgtaacaactccgccccattgacgcaaatgggcggtaggcgtgtacggtggg
aggtctatataagcagagctctctggctaactaccggtgccaccatggcccaaagaagaagcggaaggtcggtatccacggagtcccagc
agccaagcggaactacatcctgggcctggacatcggcatcaccagcgtgggctacggcatcatcgactacgagacacgggacgtgatcg
atgccggcgtgcggctgttcaaagaggccaacgtggaaaacaacgagggcaggcggagcaagagaggcgccagaaggctgaagcgg
cggaggcggcatagaatccagagagtgaagaagctgctgttcgactacaacctgctgaccgaccacagcgagctgagcggcatcaaccc
ctacgaggccagagtgaagggcctgagccagaagctgagcgaggaagagttctctgccgccctgctgcacctggccaagagaagaggc
gtgcacaacgtgaacgaggtggaagaggacaccggcaacgagctgtccaccaaagagcagatcagccggaacagcaaggccctgga
agagaaatacgtggccgaactgcagctggaacggctgaagaaagacggcgaagtgcggggcagcatcaacagattcaagaccagcga
ctacgtgaaagaagccaaacagctgctgaaggtgcagaaggcctaccaccagctggaccagagcttcatcgacacctacatcgacctgct
ggaaacccggcggacctactatgagggacctggcgagggcagccccttcggctggaaggacatcaaagaatggtacgagatgctgatg
ggccactgcacctacttccccgaggaactgcggagcgtgaagtacgcctacaacgccgacctgtacaacgccctgaacgacctgaacaat
ctcgtgatcaccagggacgagaacgagaagctggaatattacgagaagttccagatcatcgagaacgtgttcaagcagaagaagaagcc
caccctgaagcagatcgccaaagaaatcctcgtgaacgaagaggatattaagggctacagagtgaccagcaccggcaagcccgagttca
ccaacctgaaggtgtaccacgacatcaaggacattaccgcccggaaagagattattgagaacgccgagctgctggatcagattgccaaga
tcctgaccatctaccagagcagcgaggacatccaggaagaactgaccaatctgaactccgagctgacccaggaagagatcgagcagatc
tctaatctgaagggctataccggcacccacaacctgagcctgaaggccatcaacctgatcctggacgagctgtggcacaccaacgacaac
cagatcgctatcttcaaccggctgaagctggtgcccaagaaggtggacctgtcccagcagaaagagatccccaccaccctggtggacgac
ttcatcctgagccccgtcgtgaagagaagcttcatccagagcatcaaagtgatcaacgccatcatcaagaagtacggcctgcccaacgaca
tcattatcgagctggcccgcgagaagaactccaaggacgcccagaaaatgatcaacgagatgcagaagcggaaccggcagaccaacga

gcggatcgaggaaatcatccggaccaccggcaaagagaacgccaagtacctgatcgagaagatcaagctgcacgacatgcaggaagg
caagtgcctgtacagcctggaagccatccctctggaagatctgctgaacaaccccttcaactatgaggtggaccacatcatccccagaagc
gtgtccttcgacaacagcttcaacaacaaggtgctcgtgaagcaggaagaaaacagcaagaagggcaaccggaccccattccagtacct
gagcagcagcgacagcaagatcagctacgaaaccttcaagaagcacatcctgaatctggccaagggcaagggcagaatcagcaagacc
aagaaagagtatctgctggaagaacgggacatcaacaggttctccgtgcagaaagacttcatcaaccggaacctggtggataccagatac
gccaccagaggcctgatgaacctgctgcggagctacttcagagtgaacaacctggacgtgaaagtgaagtccatcaatggcggcttcacc
agctttctgcggcggaagtggaagtttaagaaagagcggaacaaggggtacaagcaccacgccgaggacgccctgatcattgccaacgc
cgatttcatcttcaaagagtggaagaaactggacaaggccaaaaaagtgatggaaaaccagatgttcgaggaaaagcaggccgagagca
tgcccgagatcgaaaccgagcaggagtacaaagagatcttcatcacccccccaccagatcaagcacattaaggacttcaaggactacaagt
acagccaccgggtggacaagaagcctaatagagagctgattaacgacaccctgtactccacccggaaggacgacaagggcaacaccct
gatcgtgaacaatctgaacggcctgtacgacaaggacaatgacaagctgaaaaagctgatcaacaagagccccgaaaagctgctgatgta
ccaccacgacccccagacctaccagaaactgaagctgattatggaacagtacggcgacgagaagaatcccctgtacaagtactacgagg
aaaccgggaactacctgaccaagtactccaaaaaggacaacggccccgtgatcaagaagattaagtattacggcaacaaactgaacgccc
atctggacatcaccgacgactaccccaacagcagaaacaaggtcgtgaagctgtccctgaagccctacagattcgacgtgtacctggaca
atggcgtgtacaagttcgtgaccgtgaagaatctggatgtgatcaaaaaagaaaactactacgaagtgaatagcaagtgctatgaggaagct
aagaagctgaagaagatcagcaaccaggccgagtttatcgcctccttctacaacaacgatctgatcaagatcaacggcgagctgtatagag
tgatcggcgtgaacaacgacctgctgaaccggatcgaagtgaacatgatcgacatcacctaccgcgagtacctggaaaacatgaacgaca
agaggcccccccaggatcattaagacaatcgcctccaagacccagagcattaagaagtacagcacagacattctgggcaacctgtatgaag
tgaaatctaagaagcaccctcagatcatcaaaaagggcaaaaggccggcggccacgaaaaaggccggccaggcaaaaaagaaaaagg
gatcctacccatacgatgttccagattacgcttacccatacgatgttccagattacgcttacccatacgatgttccagattacgcttaagaattcc
tagagctcgctgatcagcctcgactgtgccttctagttgccagccatctgttgtttgcccctcccccgtgccttccttgaccctggaaggtgcca
ctcccactgtcctttcctaataaaatgaggaaattgcatcgcattgtctgagtaggtgtcattctattctggggggtggggtggggcaggacag
caagggggaggattgggaagagaatagcaggcatgctggggaggtaccgagggcctatttcccatgattccttcatatttgcatatacgata
caaggctgttagagagataattggaattaatttgactgtaaacacaaagatattagtacaaaatacgtgacgtagaaagtaataatttcttgggt
agtttgcagttttaaaattatgttttaaaatggactatcatagaaacaccggagaccacggcaggtctcaagttttagtactctggaaacagaat
ctactaaaacaaggcaaaatgccgtgtttatctcgtcaacttgttggcgaga

**PX601-Sacas9-hpfh1: CMV-NLS-SaCas9-NLS-3xHA; U6-hpfh1-SagRNA**

CMV promoters colored in green, human codon optimized S. aureus Cas9 colored in blue, NLS underlined, U6 promoters colored in yellow, hpfh1-SagRNA colored in purple.

ggtgatgcggttttggcagtacatcaatgggcgtggatagcggtttgactcacggggatttccaagtctccaccccattgacgtcaatgggag
tttgttttggcaccaaaatcaacgggactttccaaaatgtcgtaacaactccgccccattgacgcaaatgggcggtaggcgtgtacggtggg

aggtctatataagcagagctctctggctaactaccggtgccaccatggcccaaagaagaagcggaaggtcggtatccacggagtcccagc
agccaagcggaactacatcctgggcctggacatcggcatcaccagcgtgggctacggcatcatcgactacgagacacgggacgtgatcg
atgccggcgtgcggctgttcaaagaggccaacgtggaaaacaacgagggcaggcggagcaagagaggcgccagaaggctgaagcgg
cggaggcggcatagaatccagagagtgaagaagctgctgttcgactacaacctgctgaccgaccacagcgagctgagcggcatcaaccc
ctacgaggccagagtgaagggcctgagccagaagctgagcgaggaagagttctctgccgccctgctgcacctggccaagagaagaggc
gtgcacaacgtgaacgaggtggaagaggacaccggcaacgagctgtccaccaaagagcagatcagccggaacagcaaggccctgga
agagaaatacgtggccgaactgcagctggaacggctgaagaaagacggcgaagtgcggggcagcatcaacagattcaagaccagcga
ctacgtgaaagaagccaaacagctgctgaaggtgcagaaggcctaccaccagctggaccagagcttcatcgacacctacatcgacctgct
ggaaacccggcggacctactatgagggacctggcgagggcagccccttcggctggaaggacatcaaagaatggtacgagatgctgatg
ggccactgcacctacttccccgaggaactgcggagcgtgaagtacgcctacaacgccgacctgtacaacgccctgaacgacctgaacaat
ctcgtgatcaccagggacgagaacgagaagctggaatattacgagaagttccagatcatcgagaacgtgttcaagcagaagaagaagcc
caccctgaagcagatcgccaaagaaatcctcgtgaacgaagaggatattaagggctacagagtgaccagcaccggcaagcccgagttca
ccaacctgaaggtgtaccacgacatcaaggacattaccgcccggaaagagattattgagaacgccgagctgctggatcagattgccaaga
tcctgaccatctaccagagcagcgaggacatccaggaagaactgaccaatctgaactccgagctgacccaggaagagatcgagcagatc
tctaatctgaagggctataccggcacccacaacctgagcctgaaggccatcaacctgatcctggacgagctgtggcacaccaacgacaac
cagatcgctatcttcaaccggctgaagctggtgcccaagaaggtggacctgtcccagcagaaagagatccccaccaccctggtggacgac
ttcatcctgagccccgtcgtgaagagaagcttcatccagagcatcaaagtgatcaacgccatcatcaagaagtacggcctgcccaacgaca
tcattatcgagctggcccgcgagaagaactccaaggacgcccagaaaatgatcaacgagatgcagaagcggaaccggcagaccaacga
gcggatcgaggaaatcatccggaccaccggcaaagagaacgccaagtacctgatcgagaagatcaagctgcacgacatgcaggaagg
caagtgcctgtacagcctggaagccatccctctggaagatctgctgaacaaccccttcaactatgaggtggaccacatcatccccagaagc
gtgtccttcgacaacagcttcaacaacaaggtgctcgtgaagcaggaagaaaacagcaagaagggcaaccggaccccattccagtacct
gagcagcagcgacagcaagatcagctacgaaaccttcaagaagcacatcctgaatctggccaagggcaagggcagaatcagcaagacc
aagaaagagtatctgctggaagaacgggacatcaacaggttctccgtgcagaaagacttcatcaaccggaacctggtggataccagatac
gccaccagaggcctgatgaacctgctgcggagctacttcagagtgaacaacctggacgtgaaagtgaagtccatcaatggcggcttcacc
agctttctgcggcggaagtggaagtttaagaaagagcggaacaaggggtacaagcaccacgccgaggacgccctgatcattgccaacgc
cgatttcatcttcaaagagtggaagaaactggacaaggccaaaaaagtgatggaaaaccagatgttcgaggaaaagcaggccgagagca
tgcccgagatcgaaaccgagcaggagtacaaagagatcttcatcacccccccaccagatcaagcacattaaggacttcaaggactacaagt
acagccaccgggtggacaagaagcctaatagagagctgattaacgacaccctgtactccacccggaaggacgacaagggcaacaccct
gatcgtgaacaatctgaacggcctgtacgacaaggacaatgacaagctgaaaaagctgatcaacaagagccccgaaaagctgctgatgta
ccaccacgacccccagacctaccagaaactgaagctgattatggaacagtacggcgacgagaagaatcccctgtacaagtactacgagg
aaaccgggaactacctgaccaagtactccaaaaaggacaacggccccgtgatcaagaagattaagtattacggcaacaaactgaacgccc
atctggacatcaccgacgactaccccaacagcagaaacaaggtcgtgaagctgtccctgaagccctacagattcgacgtgtacctggaca
atggcgtgtacaagttcgtgaccgtgaagaatctggatgtgatcaaaaaagaaaactactacgaagtgaatagcaagtgctatgaggaagct
aagaagctgaagaagatcagcaaccaggccgagtttatcgcctccttctacaacaacgatctgatcaagatcaacggcgagctgtatagag
tgatcggcgtgaacaacgacctgctgaaccggatcgaagtgaacatgatcgacatcacctaccgcgagtacctggaaaacatgaacgaca
agaggccccccaggatcattaagacaatcgcctccaagacccagagcattaagaagtacagcacagacattctgggcaacctgtatgaag

tgaaatctaagaagcaccctcagatcatcaaaaagggcaaaaggccggcggccacgaaaaaggccggccaggcaaaaaagaaaaagg
gatcctacccatacgatgttccagattacgcttacccatacgatgttccagattacgcttacccatacgatgttccagattacgcttaagaattcc
tagagctcgctgatcagcctcgactgtgccttctagttgccagccatctgttgtttgcccctcccccgtgccttccttgaccctggaaggtgcca
ctcccactgtcctttcctaataaaatgaggaaattgcatcgcattgtctgagtaggtgtcattctattctggggggtggggtggggcaggacag
caagggggaggattgggaagagaatagcaggcatgctggggaggtaccgagggcctatttcccatgattccttcatatttgcatatacgata
caaggctgttagagagataattggaattaatttgactgtaaacacaaagatattagtacaaaatacgtgacgtagaaagtaataatttcttgggt
agtttgcagttttaaaattatgttttaaaatggactatcatagaaacaccgttgggagatgccataaagcacaagttttagtactctggaaacaga
atctactaaaacaaggcaaaatgccgtgtttatctcgtcaacttgttggcgaga

**PX601-SaHycas9-hpfh1: CMV-NLS-SaHycas9-NLS-3xHA; U6-hpfh1- AagRNA**

CMV promoters colored in green, human codon optimized **SaHycas9** colored in blue, NLS underlined, U6 promoters colored in yellow, hpfh1-SagRNA colored in purple.

ggtgatgcggttttggcagtacatcaatgggcgtggatagcggtttgactcacggggatttccaagtctccaccccattgacgtcaatgggag
tttgttttggcaccaaaatcaacgggactttccaaaatgtcgtaacaactccgccccattgacgcaaatgggcggtaggcgtgtacggtggg
aggtctatataagcagagctctctggctaactaccggtgccaccatggcccaaagaagaagcggaaggtcggtatccacggagtcccagc
agccaagcggaactacatcctgggcctggacatcggcatcaccagcgtgggctacggcatcatcgactacgagacacgggacgtgatcg
atgccggcgtgcggctgttcaaagaggccaacgtggaaaacaacgagggcaggcggagcaagagaggcgccagaaggctgaagcgg
cggaggcggcatagaatccagagagtgaagaagctgctgttcgactacaacctgctgaccgaccacagcgagctgagcggcatcaaccc
ctacgaggccagagtgaagggcctgagccagaagctgagcgaggaagagttctctgccgccctgctgcacctggccaagagaagaggc
gtgcacaacgtgaacgaggtggaagaggacaccggcaacgagctgtccaccaaagagcagatcagccggaacagcaaggccctgga
agagaaatacgtggccgaactgcagctggaacggctgaagaaagacggcgaagtgcggggcagcatcaacagattcaagaccagcga
ctacgtgaaagaagccaaacagctgctgaaggtgcagaaggcctaccaccagctggaccagagcttcatcgacacctacatcgacctgct
ggaaacccggcggacctactatgagggacctggcgagggcagccccttcggctggaaggacatcaaagaatggtacgagatgctgatg
ggccactgcacctacttccccgaggaactgcggagcgtgaagtacgcctacaacgccgacctgtacaacgccctgaacgacctgaacaat
ctcgtgatcaccagggacgagaacgagaagctggaatattacgagaagttccagatcatcgagaacgtgttcaagcagaagaagaagcc
caccctgaagcagatcgccaaagaaatcctcgtgaacgaagaggatattaagggctacagagtgaccagcaccggcaagcccgagttca
ccaacctgaaggtgtaccacgacatcaaggacattaccgcccggaaagagattattgagaacgccgagctgctggatcagattgccaaga
tcctgaccatctaccagagcagcgaggacatccaggaagaactgaccaatctgaactccgagctgacccaggaagagatcgagcagatc
tctaatctgaagggctataccggcacccacaacctgagcctgaaggccatcaacctgatcctggacgagctgtggcacaccaacgacaac
cagatcgctatcttcaaccggctgaagctggtgcccaagaaggtggacctgtcccagcagaaagagatccccaccaccctggtggacgac
ttcatcctgagccccgtcgtgaagagaagcttcatccagagcatcaaagtgatcaacgccatcatcaagaagtacggcctgcccaacgaca
tcattatcgagctggcccgcgagaagaactccaaggacgcccagaaaatgatcaacgagatgcagaagcggaaccggcagaccaacga

gcggatcgaggaaatcatccggaccaccggcaaagagaacgccaagtacctgatcgagaagatcaagctgcacgacatgcaggaagg
caagtgcctgtacagcctggaagccatccctctggaagatctgctgaacaacccctttcaactatgaggtggaccacatcatccccagaagc
gtgtccttcgacaacagcttcaacaacaaggtgctcgtgaagcaggaagaaaacagcaagaagggcaaccggaccccattccagtacct
gagcagcagcgacagcaagatcagctacgaaaccttcaagaagcacatcctgaatctggccaagggcaagggcagaatcagcaagacc
aagaaagagtatctgctggaagaacgggacatcaacaggttctccgtgcagaaagacttcatcaaccggaacctggtggataccagatac
gccaccagaggcctgatgaacctgctgcggagctacttcagagtgaacaacctggacgtgaaagtgaagtccatcaatggcggcttcacc
agctttctgcggcggaagtggaagtttaagaaagagcggaacaaggggtacaagcaccacgccgaggacgccctgatcattgccaacgc
cgatttcatcttcaaagagtggaagaaactggacaaggccaaaaaagtgatggaaaaccagatgttcgaggaaaagcaggccgagagca
tgcccgagatcgaaaccgagcaggagtacaaagagatcttcatcaccccccaccagatcaagcacattaaggacttcaaggactacaagt
acagccacagagtggacaagaagccaaaccgccagctgattaacgacacactgtactccacccggaacattgacggcattgagtacgtg
gtggagtctattaaggacatttactccgtgaacaacgacaaggtaaaaacaaagttcaagaaggaccctcacaggctgctcatgtaccgga
acgaccctcagaccttcgagaagttcgagaaggtgttcaagcagtacgagtctgagaagaaccctttcgccaagtactacgaggagacag
gcgagaagattcgcaagttctctaagacaggccagggcccatacatcaacaagattaagtacctgcgggagcgcctgggccggcactgc
gacgtgaccaacaagtacatcaactctcggaacaagatcgtgcagctgaagatttactcttaccgcttcgacatttaccagtacggcaacaac
tacaagatgatcaccatcagctacattgacctggagcagaagtctaactactactacatttctagggagaagtacgagcagaagaagaagg
acaagcagatcgacgactcttacaagttcattggctccttctacaagaacgacattattaactacaacggcgagatgtaccgcgtgattggcg
tgaacgactctgagaagaacaagatccagctggacatgattgacatctctattaaggactacatggagctgaacaacattaaaaaaaccggc
gtgatttacaagaccatcggcaagtccaccacacacatcgagaagtacaccacagacatcctgggcaacctgtacaaggccgcccctcct
aagaagcctcagctcatcttcaagaaaaggccggcggccacgaaaaaggccggccaggcaaaaaagaaaaagggatcctacccatacg
atgttccagattacgcttacccatacgatgttccagattacgcttacccatacgatgttccagattacgcttaagaattcctagagctcgctgatc
agcctcgactgtgccttctagttgccagccatctgttgtttgcccctcccccgtgccttccttgaccctggaaggtgccactcccactgtcctttc
ctaataaaatgaggaaattgcatcgcattgtctgagtaggtgtcattctattctggggggtggggtggggcaggacagcaagggggaggatt
gggaagagaatagcaggcatgctggggaggtaccgagggcctatttcccatgattccttcatatttgcatatacgatacaaggctgttagaga
gataattggaattaatttgactgtaaacacaaagatattagtacaaaatacgtgacgtagaaagtaataatttcttgggtagtttgcagttttaaaa
ttatgttttaaaatggactatcatagaaacaccgttgggagatgccataaagcacaagttttagtactctggaaacagaatctactaaaacaag
gcaaaatgccgtgtttatctcgtcaacttgttggcgaga

**pLV-NNGRRT: CMV-hpfh1-NNGRRT-GFP**

CMV promoters colored in green, hpfh1 target sequence colored in blue, NNGRRT underlined, GFP colored in purple.

cgttacataacttacggtaaatggcccgcctggctgaccgcccaacgacccccgcccattgacgtcaataatgacgtatgttcccatagtaac
gccaatagggactttccattgacgtcaatgggtggagtatttacggtaaactgcccacttggcagtacatcaagtgtatcatatgccaagtacg

ccccctattgacgtcaatgacggtaaatggcccgcctggcattatgcccagtacatgaccttatgggactttcctacttggcagtacatctacgt
attagtcatcgctattaccatggtgatgcggttttggcagtacatcaatgggcgtggatagcggtttgactcacggggatttccaagtctccacc
ccattgacgtcaatgggagtttgttttggcaccaaaatcaacgggactttccaaaatgtcgtaacaactccgccccattgacgcaaatgggcg
gtaggcgtgtacggtgggaggtctatataagcagagctttctagaaatgtacaaggaattcatggaacgttgggagatgccataaagcacc
tggatgtgagcaagggcgaggagctgttcaccggggtggtgcccatcctggtcgagctggacggcgacgtaaacggccacaagttcagc
gtgtccggcgagggcgagggcgatgccacctacggcaagctgaccctgaagttcatctgcaccaccggcaagctgcccgtgccctggcc
caccctcgtgaccaccctgacctacggcgtgcagtgcttcagccgctaccccgaccacatgaagcagcacgacttcttcaagtccgccatg
cccgaaggctacgtccaggagcgcaccatcttcttcaaggacgacggcaactacaagacccgcgccgaggtgaagttcgagggcgaca
ccctggtgaaccgcatcgagctgaagggcatcgacttcaaggaggacggcaacatcctggggcacaagctggagtacaactacaacagc
cacaacgtctatatcatggccgacaagcagaagaacggcatcaaggtgaacttcaagatccgccacaacatcgaggacggcagcgtgca
gctcgccgaccactaccagcagaacacccccatcggcgacggccccgtgctgctgcccgacaaccactacctgagcacccagtccgcc
ctgagcaaagaccccaacgagaagcgcgatcacatggtcctgctggagttcgtgaccgccgccgggatcactctcggcatggacgagct
gtacaagtaaggatcc

**pLV-NNAAAA: CMV-hpfh1-NNAAAA-GFP**

CMV promoters colored in green, hpfh1 target sequence colored in blue, NNGRRT underlined, GFP colored in purple.

cgttacataacttacggtaaatggcccgcctggctgaccgcccaacgacccccgcccattgacgtcaataatgacgtatgttcccatagtaac
gccaatagggactttccattgacgtcaatgggtggagtatttacggtaaactgcccacttggcagtacatcaagtgtatcatatgccaagtacg
ccccctattgacgtcaatgacggtaaatggcccgcctggcattatgcccagtacatgaccttatgggactttcctacttggcagtacatctacgt
attagtcatcgctattaccatggtgatgcggttttggcagtacatcaatgggcgtggatagcggtttgactcacggggatttccaagtctccacc
ccattgacgtcaatgggagtttgttttggcaccaaaatcaacgggactttccaaaatgtcgtaacaactccgccccattgacgcaaatgggcg
gtaggcgtgtacggtgggaggtctatataagcagagctttctagaaatgtacaaggaattcatggaacgttgggagatgccataaagcacc
taaaagtgagcaagggcgaggagctgttcaccggggtggtgcccatcctggtcgagctggacggcgacgtaaacggccacaagttcagc
gtgtccggcgagggcgagggcgatgccacctacggcaagctgaccctgaagttcatctgcaccaccggcaagctgcccgtgccctggcc
caccctcgtgaccaccctgacctacggcgtgcagtgcttcagccgctaccccgaccacatgaagcagcacgacttcttcaagtccgccatg
cccgaaggctacgtccaggagcgcaccatcttcttcaaggacgacggcaactacaagacccgcgccgaggtgaagttcgagggcgaca
ccctggtgaaccgcatcgagctgaagggcatcgacttcaaggaggacggcaacatcctggggcacaagctggagtacaactacaacagc
cacaacgtctatatcatggccgacaagcagaagaacggcatcaaggtgaacttcaag~~at~~ccgccacaacatcgaggacggcagcgtgca
gctcgccgaccactaccagcagaacacccccatcggcgacggccccgtgctgctgcccgacaaccactacctgagcacccagtccgcc

ctgagcaaagaccccaacgagaagcgcgatcacatggtcctgctggagttcgtgaccgccgccgggatcactctcggcatggacgagct
gtacaagtaaggatcc

**DNA Targets for PAM Specificity (Figure 5)**

| Gene | Spacer | 5'-PAM-3' |
|---|---|---|
| hpfh1 | TTGGGAGATGCCATAAAGCAC | NNGGAT(NNGRRT) |
| hpfh1 | TTGGGAGATGCCATAAAGCAC | NNAAAA |

**T7E1 Primers**

| Name | Forward | Reverse | Tm (°C) |
|---|---|---|---|
| hpfh1 outer prime | TACATGACCTTATGGGACTTT CCT ACTTGGCAGTA | GCCAGATATAGACGTTGTG GCTGTTGTAGTTGTA | 71 |
| hpfh1 inner prime | GGATAGCGGTTTGACTCACGG | CAGGATGTTGCCGTCCTCCTT | 65 |
